# HPV-positive head and neck squamous cell carcinoma cells lose viability during triggered myocyte lineage differentiation

**DOI:** 10.1101/2024.03.28.587125

**Authors:** Sarah Gendreizig, Laura Martinez-Ruiz, Alba López-Rodríguez, Harkiren Pabla, Leonie Hose, Frank Brasch, Tobias Busche, Germaine Escames, Holger Sudhoff, Lars Uwe Scholtz, Ingo Todt, Felix Oppel

## Abstract

Head and neck squamous cell carcinoma (HNSCC) is a highly malignant disease, and death rates have remained at approximately 50% for decades. New tumor-targeting treatment strategies are desperately needed. Using patient-derived tumor cells, we created an HNSCC differentiation model of HPV+ tumor cells from two patients. We observed a loss of malignant characteristics in differentiating cell culture conditions, including irregularly enlarged cell morphology, cell cycle arrest with downregulation of Ki67, and reduced cell viability. RNA-seq showed myocyte-like differentiation with upregulation of markers of myofibril assembly, including TPM1, TAGLN, and ACTA1. Immunofluorescence staining of differentiated and undifferentiated primary HPV+ HNSCC cells confirmed an upregulation of these markers and the formation of parallel actin fibers reminiscent of myoblast-lineage cells. Moreover, immunofluorescence of HPV+ tumor tissue revealed areas of cells co-expressing the identified markers of myofibril assembly, HPV surrogate marker p16, and stress-associated basal keratinocyte marker KRT17, indicating that the observed myocyte-like *in vitro* differentiation occurs in human tissue. A recent sarcoma study was able to turn rhabdomyosarcoma into muscle-like cells. We are the first to report that carcinoma cells can undergo a triggered myocyte differentiation. Our study suggests that the targeted myo-differentiation of tumor cells might be therapeutically valuable in HPV+ HNSCCs.

## Background

Head and neck squamous cell carcinomas (HNSCC) are a diverse group of cancers originating from various head and neck regions, including the nasopharynx, oral cavity, oropharynx, hypopharynx, and larynx. Globally, HNSCC accounts for roughly 700,000 new cases and 380,000 deaths annually (1). The 3- and 5-year overall survival rates for human papillomaviruses (HPV) positive HNSCC patients are 80.0% and 75.0%, and for HPV-negative patients, 54.0% and 48.0% (2). Due to this reason, it is imperative to develop new treatment strategies that specifically target tumors. The human papillomaviruses are a group of common viruses that infect skin and mucous membranes. They can cause warts, which usually go away on their own, but certain types of HPV can cause cancer. There are over 100 types of HPV, which are classified into two groups: low-risk and high-risk (oncogenic). The low-risk HPVs, mostly HPV6 and 11, can cause harmless and temporary lesions. However, high-risk HPV infection may lead to malignant transformation. The high-risk group includes HPV16, 18, 31, 33, 35, 39, 45, 51, 52, 56, 58, 59, and 68 (3).

Differentiation therapy (DTH) is a therapeutic strategy that involves the utilization of diverse molecular agents capable of inducing differentiation in malignant cells. The rationale behind this approach is to promote the differentiation of cancer cells into specialized cell types, resulting in the removal of these cells from the proliferative compartment. This approach aims to promote terminal differentiation, which is regarded as a final branch of cellular development. Consequently, DTH has emerged as a promising treatment modality for cancer and has garnered significant attention in the scientific community (4).

A previous study discovered that HPV-negative HNSCC cell differentiation can be induced through cornification, leading to a loss of cell malignancy (5). This finding introduced a new strategy and opportunity for targeted therapy in HPV-negative related HNSCC. The successful reprogramming of cancer cells into differentiated cells has been implemented before, most notably in PML-RARα fusion-driven acute promyelocytic leukemia (6). Nevertheless, it remains a challenge in other types of tumors, such as colon cancer (COCA) and pancreatic ductal adenocarcinoma (PDAC). The reason is that tumor-initiating cells (TICs) possess the ability to change their characteristics and functions, allowing tumors to avoid terminal differentiation through epigenetic mechanisms ( (7), (8), (9)). This phenomenon resembles the self-renewing properties of normal tissue regeneration (10). Unfortunately, this characteristic makes these types of cancers unsuitable for tumor differentiation therapies to date. However, our previous study found that differentiation therapy might represent an effective treatment strategy for HPV-negative HNSCCs. This is because terminal cornification, a non-reversible process, leads to programmed cell death and loss of malignant behavior. Indicating that cancer that arises from skin or mucosal tissue undergoing natural cornification appears to be vulnerable to this type of therapy. Further research is required to prove this point.

The number of HPV+ immortalized HNSCC cell lines available is significantly limited; precisely eight lines are known ( (11), (12), (13)). Developing primary cell cultures from naturally infected HPV-positive cancers is a challenging task (14). However, we have managed to provide two such cell culture models. As of now, there is no existing documentation for a primary HPV-positive HNSCC differentiation model. Our system represents an opportunity to learn more about differentiation in this tumor type and may aid the identification of new therapy targets to target HPV-associated cancer types. This report provides an overview of the establishment and characterization of two *in vitro* HPV-positive HNSCC differentiation models and discusses their potential for cancer therapy.

## Materials and methods

### Human material and cell culture

Primary head and neck cancer tissue was obtained from medically indicated surgeries with informed consent of the patients, according to the declaration of Helsinki, and as approved by the ethics committee of the Ruhr-University Bochum (AZ 2018-397_3), as reported previously (14). The cells were cultured in PneumaCult™-Ex Plus Medium, referred to as stem cell medium (SCM, #05041, Stemcell Technologies, Vancouver, Canada) supplemented as described (15) Adherent cells and spheroids were detached using Accutase (Capricorn Scientific, Ebsdorfergrund, Germany). Cells were differentiated in cardiac fibroblast medium (CFM, 316K-500, Cell Applications, San Diego, CA, United States) supplemented with 1% 200 mM L-Glutamine, 1% 100x Penicillin/Streptomycin, and 1% 250 μg/mL Amphotericin B solution (all three Capricorn Scientific). Dulbecco’s Modified Eagle Medium (DMEM, DMEM-HXA, Capricorn Scientific) was supplemented with 10% Fetal Bovine Serum Advanced (FBS Advanced, FBS-11A, Capricorn Scientific), L-Glutamine, Penicillin/Streptomycin, and Amphotericin B. Cells received fresh medium twice a week. Cells were analyzed for proliferation and cornification and treated with drugs and cytokines as outlined in supplementary methods.

### Mouse xenograft tumor models

All animal experiments were conducted as approved by the Institutional Animal Care and Use Committee of the University of Granada (procedures 12/07/2019/128) in accordance with the European Convention for the Protection of Vertebrate Animals used for Experimental and Other Scientific Purposes (CETS #123) and Spanish law (R.D. 53/2013).

NSG mice (NOD.Cg-Prkdcscid Il2rgtm1Wjl/SzJ, The Jackson Laboratory, Bar Harbor, ME, United States) were housed in appropriate sterile filter-capped cages with sterile food and water ad libitum. 1x10^6^ cells were transplanted subcutaneously as described previously (14), with differentiated and undifferentiated cell populations being injected side-by-side into the left and right flank of each mouse as performed in a previous study (8). Animals were monitored for tumor development twice a week. Tumor sizes were measured using a vernier caliper and tumor volume was calculated by the formula (width×length2)/2. When the first tumor of each population reached the legal size limit, all mice of that experiment were measured and sacrificed. Tumors were harvested and fixed in 4% paraformaldehyde (PFA, 2.529.311.214, PanReac) for 24 hours. The fixed samples were embedded in paraffin following standard protocols.

### Histopathology analysis of HNSCC tissue

Paraffin-embedded tissue was sectioned with 2 μm thickness using a sliding HM430 microtome (Zeiss). Hematoxylin/Eosin (HE) staining was performed using standard protocols in a linear COT 20 tissue stainer (MEDITE, Burgdorf, Germany). HE stained section were analyze by a senior pathologist (F.B.) to compare patient characteristics between original tumor and xenograft tumor models. To quantify keratinized areas in human tumor tissue every HE-stained section was subdivided into three random views and analyzed using an Axio Lab.1 microscope (Zeiss) and DISKUS software (Hilgers, Königswinter, Germany). Per view, the total area displaying malignant histology was determined in mm^2^. Next, the keratinized area within the tumor compartment was determined to calculate the proportional extent of cornification using Microsoft Excel.

### Cornification Assay

Cornification was quantified based on a previous protocol to isolate cornified envelopes (16). Triplicates of P1 and P2 cells were seeded into 6-well plates in SCM or CFM until cells were 80-90% confluent. Cells were washed with 1x PBS, detached using Accutase (Capricorn Scientific, Ebsdorfergrund, Germany), and stained with trypan blue (Sigma Aldrich), followed by a count of live and dead cells using a Neubauer Chamber. The cells were washed in 1x PBS and subsequently mixed with 100 μL 4% SDS (Carl Roth GmbH, Karlsruhe, Germany) and 2% beta-Mercaptoethanol (Merck, Darmstadt, Germany) in PBS. The suspension is cooked at 95°C for 5-10 min. Lastly, the cornified envelopes were counted using a Neubauer Chamber.

### Proliferation Assay

Triplicates of P1 and P2 cells (5x104) were seeded into a 12-well plate in SCM or CFM. After 5 days of incubation at 37°C with 5% CO2, cells were washed with 1x PBS and detached using Accutase (Capricorn Scientific, Ebsdorfergrund, Germany). The cells were manually counted using a Neubauer Chamber. After that, cells were washed in 1x PBS, resuspended in a culture medium, and further incubated at the same conditions. The growth curves were generated over a period of 15 days, with repeated counting every 5 days.

### Indirect Immunofluorescence of HNSCC cells and tissue

Indirect Immunofluorescence (IF) of HNSCC cell cultures was performed as described previously by (17). Tumor tissue of patients and xenograft tumor models was analyzed by IF as established in prior studies (8). Samples were imaged using a LSM780 confocal microscope (Zeiss, Oberkochen, Germany) and ZEN software (Zeiss). DNA was stained with DAPI (Sigma Aldrich, St. Louis, MO, United States) using 2μg/ml diluted in PBS + 0.1% bovine serum albumin (Capricorn Scientific). Primary antibodies: rabbit-anti-human ACTA1 (1:200, EPR16769, ab179467, Abcam, Cambridge, UK), rabbit-anti-human CDKN2A/p16INK4a (EPR1473; 1:200; Abcam), rabbit-anti-human EpCam (EGP40/1556R; 1:100; Novus Biologicals, Centennial, CO, United States), mouse HPV antibody cocktail (1:100, CAMVR-1 & C1P5, Z2657MS, Thermo Fisher Scientific, Waltham, MA, United States), rabbit-anti-human Ki67 (1:200, SP6, ab16667, Abcam), mouse anti-human Tropomyosin (1:100, F-6, sc-74480, Santa Cruz Biotechology, Dallas, TX, United States). Secondary antibodies: goat-anti-mouse-IgG-Alexa Fluor-555 (1:400, A21422), donkey-anti-rabbit-IgG-Alexa Fluor-488 (1:400, A11008), and goat-anti-guinea pig-IgG-Alexa Fluor-647 (1:400, A21450, all Thermo Fisher Scientific, Waltham, MA, United States). The actin cytoskeleton was stained using phalliodin-PF647 for 30 min (1:50, Promokine, Heidelberg, Germany).

### RNA Isolation

Cells were cultured for eight days. Total RNA was isolated from cell culture (n=3) using innuPREP DNA/RNA Mini Kit (Analytik Jena, Jena, Germany) as the manufacturer’s protocol recommended. RNA quality was determined with a BioDrop Duo+ spectral photometer (Biochrom, Holliston, USA).

### Analysis of RNA-seq Data

RNA-seq analysis was performed as previously described (5).

## RESULTS

### Primary HPV-positive HNSCC cell culture model

We conducted a study to examine the effect of a potential differentiation treatment approach for HPV-positive HNSCC. For this purpose, we established two primary cell culture models, S12 and S82, derived from oropharyngeal HNSCC of males aged 49 and 81 who had tested positive for human papillomavirus 16 (Figure 1A). S12 is derived from a previously established xenograft tumor model (14). We refer to S12 as patient 1 (P1) and S82 as patient 2 (P2). P1 is a primary cell culture derived from a xenograft tumor model, while P2 was directly established from the patient’s tissue. The xenograft tumors of P1 closely resembled the original tumor in histology (Shao and Scholtz et al.), and we isolated cells from the xenograft tumor tissue that could be expanded for more than 20 passages in stem cell medium (SCM) without phenotypic changes (Figure 1B). The cells obtained from P2 could be cultured in SCM for 6 to 8 passages (Figure 1B).

**Figure 1:**
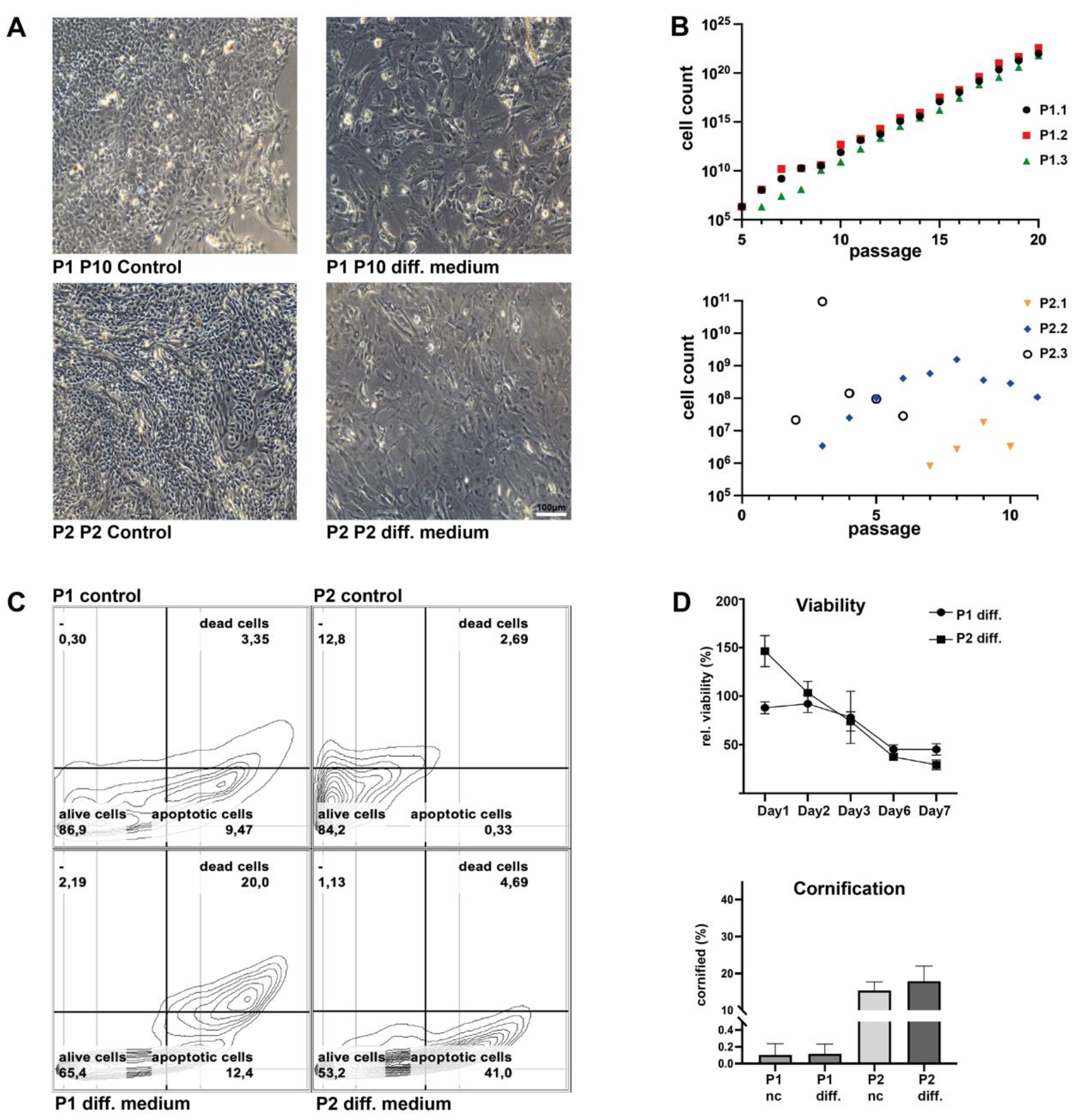
HPV+ HNSCC differentiation model reveals loss of cell viability. **(A)** Cultures of P1 and P2 display adherent monolayer cells in SCM with epithelial morphology (undifferentiated). Treatment of SCM cultures with differentiation medium (CFM) results in the loss of epithelial morphology and enlarged cells with irregular shapes. **(B)** Graphs show exponential proliferation of P1 cells for 20 passages (5 months), whereas cells of P2 stop proliferation within 11 passages; 3 independent experiments for each patient are displayed. **(C)** Apoptosis and death rate of CFM-treated cells within 72 hours; measured by Annexin-PI staining. Three technical replicates were used (n=3). **(D)** Differentiation medium treatment for 7 days reduced cell viability. Differentiation medium treatment does not increase cornification in cultures of P1 and P2. Cornification was very strong in P2 cells in both media, whereas P1 explants showed low cornification levels.

### Triggered differentiation of HPV-positive HNSCC cells activates myocyte-like gene expression

To trigger the differentiation of HPV+ HNSCC cells, we treated the cells of P1 and P2 with a differentiation medium called cardiac fibroblast medium (CFM), previously found to trigger differentiation in HPV-negative HNSCC cells (source). CFM treatment resulted in an enlarged irregular cell morphology (Figure 1A). After 72 hours of treatment, the proportion of apoptotic cells increased from 9.47% to 12.4%, and death rates were elevated by 16.65%. Meanwhile, the effect in P2 cells was delayed, with 41% of apoptotic cells and 2.69% of dead cells (Figure 1C). During the experiment, the cell viability was assessed for seven days. The results show a reduction of about 50% when compared to the SCM cultivation (Figure 1D). The cornification of HPV-positive HNSCC cells was unaffected by the differentiation treatment (Figure 1D), and there was no increase in migration (suppl. Figure 1).

RNA-seq was obtained to understand the signaling processes that lead to an increase in the death rate of our primary cell cultures due to the differentiation medium. We compared the impact of the differentiation medium on the expression of HPV-negative cells from a recent study and our HPV-positive cell cultures (Figure 2A). Our findings suggest that the differentiation medium affects both HNSCC subtypes, but the underlying mechanisms differ. Our data show 1244 differentially upregulated and 1197 downregulated genes (Figure 2A). Genes commonly upregulated upon differentiation of both HPV-positive cell cultures were ACTA1, ANKRD1, FILIPIL, FN1, PSMA6, TAGLN, and TMP1 (Figure 2B). Gene Ontology analysis of these genes revealed the activation of a myocyte-like expression pattern (Figure 2C). Significantly enriched gene sets that differentiated cells of both patients have in common include actomyosin structure organization, cellular component assembly involved in morphogenesis, muscle organ development, striated muscle cell development, and myofibril assembly (suppl. Table XX). The genes downregulated in both patients were ALDH7A1, FXYD3, HYOU1, LUC7L3, PKM, PWWP2A, SRRM2, TCEA1, and TPRKB (Figure 2B) with no significantly enriched gene sets. Differentiated cells of both patients displayed changes in their actin filament structure, shifting from a cortical distribution to a parallel fiber arrangement that resembled a myocyte phenotype. This was indicated by a strong TPM1 protein signal in indirect immunofluorescence staining (IF) that aligned with the parallel ACTIN filaments (Figure 3).

**Figure 2:**
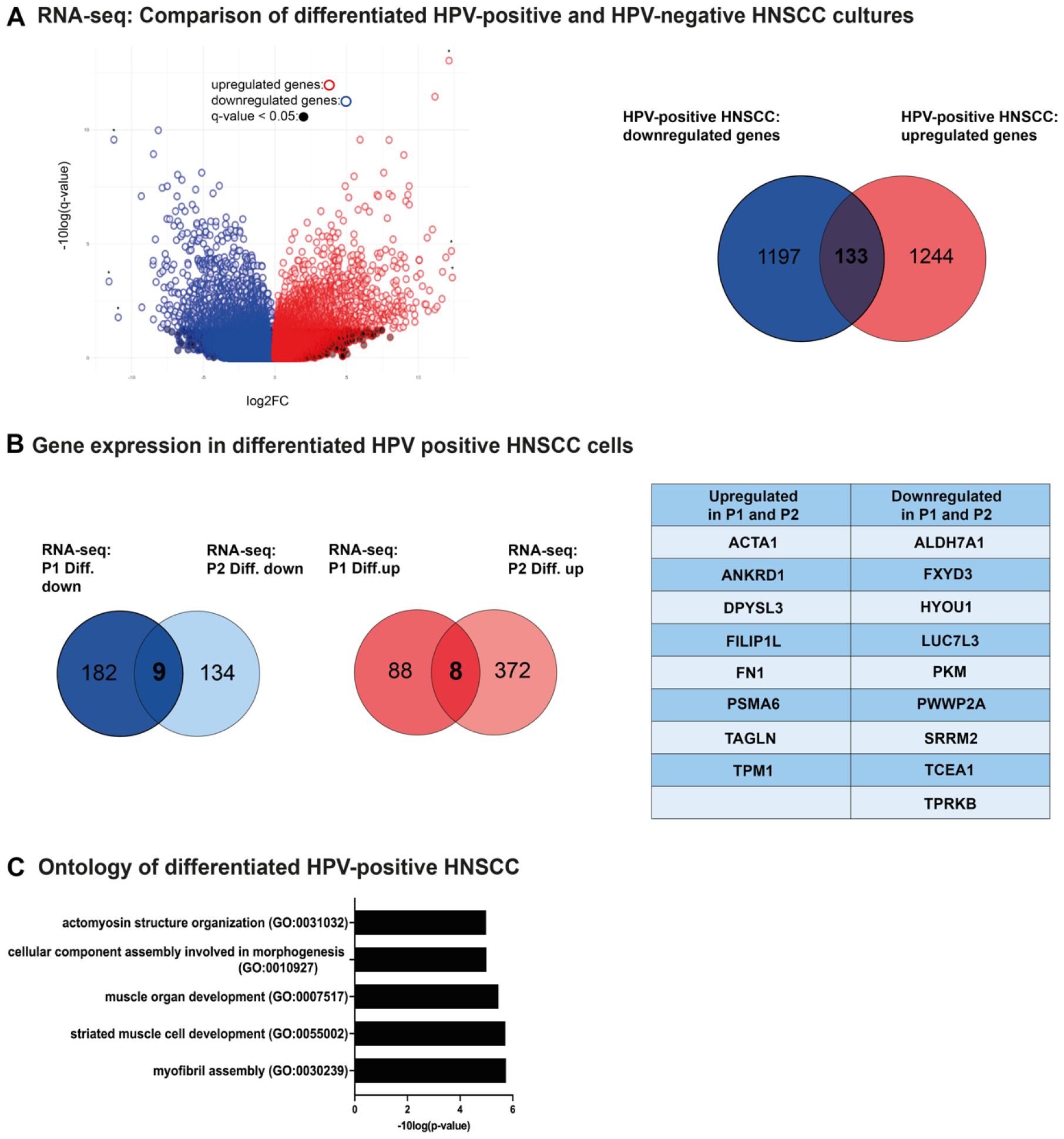
HPV-positive HNSCC cells differentiate into myocyte-like cells. **(A)** The global expression data highlights the contrasting characteristics of HPV-positive and HPV-negative HNSCC differentiation. Revealing that the mechanism of differentiation differs. **(B)** Gene expression analysis of P1 and P2 cell cultures shows either upregulated or downregulated genes upon differentiation. **(C)** Analysis of the nine genes upregulated in differentiated cells of both patients revealed five significant Gene Ontology Biological Process pathways associated with HPV+ HNSCC cell differentiation. Three technical replicates were used (n=3) with a statistical cut-off p-value of 0.05.

**Figure 3:**
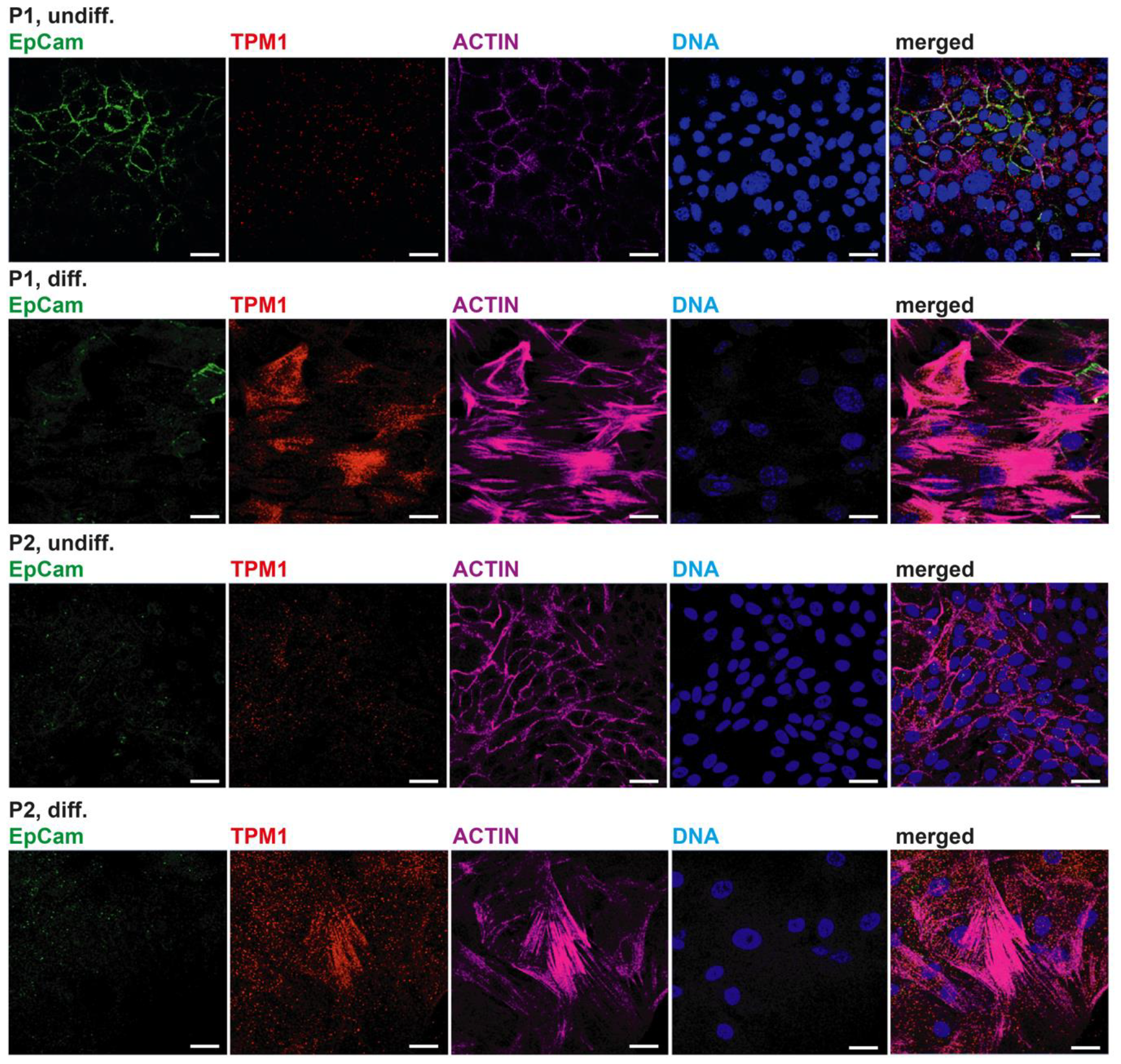
Differentiated HPV+ HNSCC cell cultures express markers of myocyte differentiation. When treated with a differentiation medium, P1 cells lose expression of EpCam, whereas cells of P2 are negative for EpCam in any medium. Both cell cultures upregulate muscle cell marker tropomyosin 1 (TPM1) and show a reorganization of the actin cytoskeleton from cortical actin to the formation of parallel fibers, reminiscent of muscle lineage differentiation; scale bars = 10 μm.

In contrast, undifferentiated cells grown in the SCM medium exhibited no TPM1 expression. SCM and CFM cells of P1 and P2 expressed HPV-related proteins, but CFM treatment strikingly diminished Ki67 proliferation marker expression (suppl. Figure S2). Notably, HPV-related proteins were detected under both conditions.

### Identified myocyte-associated markers are expressed in HPV-positive HNSCC tissue

To examine the presence of myocyte markers in human tumors, an IF staining was carried out to detect TPM1, p16, and KRT17. KRT17 is a stress-keratin and an early differentiation marker in HPV-HNSCCs (5). The tumor tissue of three patients (P1, P2, and P3) and the xenograft tumor tissue of P1 from a previous study were examined. The IF revealed the abundance of cells expressing the aforementioned myocyte markers in the tumor tissues of the selected patients. We observed strong TPM1 expression in desmoplastic stromal areas and in p16/KRT17+ cells with epithelial morphology (Suppl. Figure S3). Moreover, ACTA1 was found to be expressed in cells staining co-positive for HPV-related proteins (suppl. Figure S4). Thus, a subpopulation of HPV+ tumor cells expresses myocyte markers. In the original tumor specimens of P1 and P2, these cells were diffusely spread throughout the tumor tissue. HNSCC xenograft tumors are known to display a lymph node metastasis-like cystic/necrotic growth pattern ( (5), (18)). Xenograft tumor tissue of P1 contained enlarged cells with pleomorphic nuclei towards the core of the inner cyst (Figure 4A). These stained strongly positive for KRT17 (Figure 4B). By confocal microscopy imaging of this area, we observed a correlation between TPM1 and KRT17 expression and Ki67 proliferating cells located in the TMP1-/KRT17-population (Figure 4C). This was also detected in the original tumor tissue of P3. Our data indicate that myocyte markers and KRT17 represent markers of differentiated non-proliferative HPV+ HNSCC cells.

**Figure 4:**
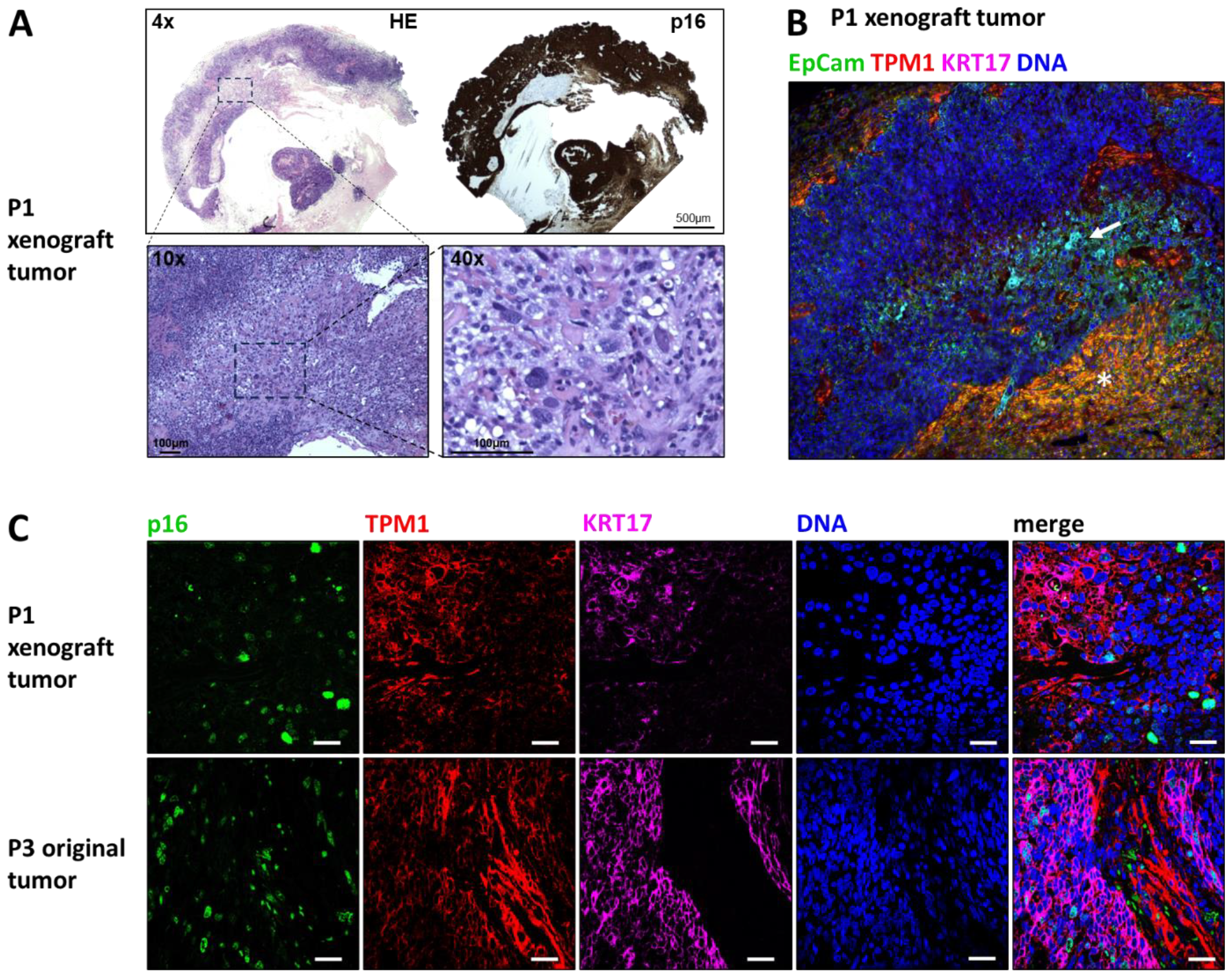
Myocyte-like differentiation in HPV+ HNSCC tissue. **(A)** Xenograft tumor of P1 with cystic/necrotic core. Hematoxylin/eosin (HE) and p16 staining of the same paraffin block were shifted into a similar orientation for comparison. **(B)** IF staining of P1 xenograft tumor reveals cells co-expressing TPM1 and KRT17 (arrow) and tissue positive for TPM1 and EpCam (star); 10x magnification. **(C)** P1 xenograft tumor and P3 original tumor show Ki67 in areas with lower TPM1/KRT17 expression; scale bars = 10 μm.

## DISCUSSION

This study successfully established two primary cell culture models of HPV-positive HNSCC. The use of primary cell cultures was imperative due to the unfavorable genetic or epigenetic changes associated with the immortalization process of cell lines. Thus, immortalized cell lines are not an adequate substitute for tumor differentiation studies (19).

Our findings demonstrate that we can force HPV-positive HNSCC cells to differentiate functionally into a robust myocyte phenotype. During this differentiation, we observed a morphological structural reorganization from an epithelial to a myocytic phenotype, indicating a successful transition. We accounted for this differentiation phenotype by demonstrating that markers like TPM1, TAGLN, and ACTA1 revealed by RNA-seq were found within both culture models. In vertebrates, TPM1 plays a crucial role in regulating striated muscle contraction that is dependent on calcium (20). TAGLN is a protein that is highly present in smooth muscle cells and is commonly used as a standard marker for these cells (21). RNA-seq revealed that actomyosin structure organization, muscle organ development, striated muscle cell differentiation, and myofibril assembly are the mechanisms behind the differentiation process. Further, the cell cultures and the original tissue strongly correlate TPM1/ACTA1 and the recently described differentiation marker KRT17 (5). Indicating that our differentiation models resemble their tissue of origin.

It was observed that different mechanisms were triggered by the same protocol used to differentiate HPV-positive and HPV-negative head and neck squamous cell carcinoma models. Our previous study showed that in HPV-negative HNSCC, the terminal differentiation was induced by cornification, resulting in the epigenetic loss of cell malignancy (5). However, our recent findings indicate that HPV-positive HNSCC cells undergo myocyte-like differentiation, which causes the loss of malignant characteristics. HPV+ HNSCC cells that have undergone induced differentiation exhibit epithelial-to-mesenchymal transition (EMT), which is a hallmark of cancer (22). Notably, in this model, the EMT transition leads to a less aggressive phenotype, resulting in reduced proliferation and viability, decreased Ki67 expression, no changes in migration, and increased apoptosis and cell death. Regarding EMT as a cancer hallmark, recent reports demonstrate that efficient invasion and metastasis require the expression of ZEB1 and SNAIL ( (23), (24)). The expression data from both of our patient-derived models did not show any evidence of such invasive or malignant signaling.

Moreover, the upregulation of Filamin A Interacting Protein 1 Like (FILIP1L) and the downregulation of ALDH7A1 induced here could be responsible for the loss of malignant characteristics. FILIP1L is described to induce loss of proliferation and migration in endothelial cells and is known to inhibit melanoma growth when expressed in tumor-associated vasculature (25). ALDH7A1 is generally known as a stemness marker in prostate cancer (26) and is involved in the growth of pancreatic ductal adenocarcinoma (27) and might thus be implicated in the maintenance of the undifferentiated phenotype of long-term repopulating HPV+ HNSCC cells. Further studies have to investigate the role of these effectors highlighted here.

The successful reprogramming of cancer cells into differentiated cells has been implemented in other malignancies, most notably PML-RARα fusion-driven acute promyelocytic leukemia (6). In the field of sarcoma, researchers at Cold Spring Harbor Laboratory have found a promising DTH approach by CRISPR-based targeting of NF-Y. The group differentiated rhabdomyosarcoma cells into muscle cells, representing that tissue’s normal differentiation path (28). To our knowledge, our team is the first to present an induced functional and morphological alteration from cancer cells into muscle-like cells in carcinomas. Both approaches differ in their experimental implementation but lead to similar results. According to our findings, the targeted induction of myogenic differentiation in tumor cells confers therapeutic benefits not only in sarcomas but also in carcinomas.

Most importantly, a clinical approach based on our data has to be established to test the suitability of our targeted differentiation for therapeutic intervention against HNSCCs. Although an *in vitro* differentiation protocol has been developed successfully here, translating it into a therapeutic drug regimen will be challenging and needs to be pursued in follow-up studies. Further research is necessary to identify an alternative composition of mediators that can mimic the impact of the CFM medium. Once a more practical treatment method is found, testing its translational value in xenograft tumor models of HNSCCs in immunodeficient mice is essential.

Our research on HPV-positive HNSCC has yielded promising results for future pharmacological studies. We observed a decrease in malignant traits by inducing myo-differentiation, which indicates that differentiation-based therapies could effectively treat this type of cancer. This method of treatment may also be useful for other HPV-associated cancers. Overall, this system could be a versatile tool for explicit pharmacological studies, leading to a more comprehensive understanding of HPV-positive HNSCC in general.

## Authors’ contributions

Conceptualization, S.G., F.O.; methodology, S.G., F.O, L.M.R. validation, S.G., F.O., and L.M.R.; formal analysis, S.G., F.O., and L.M.R.; investigation, S.G., F.O., L.M.R., A.L.R., L.H., H.P., and F.B.; resources, L.U.S., F.B., G.E., T.B., IT., and H.S.; data curation, S.G and F.O.; software, S.G.; writing—original draft preparation, S.G.; writing-review and editing, S.G., F.O., L.M.R., A.L.R., L.H., H.P., G.E., F.B., and T.B., L.U.S., I.T., and H.S.; visualization, S.G. and F.O., supervision, S.G., F.O., G.E., T.B., I.T., and H.S.; project administration, S.G., F.O., L.U.S., G.E., F.B., T.B., I.T., and H.S.; funding acquisition, F.O., G.E. and H.S..

## Competing interests

The authors declare no conflict of interest.

## Funding

This project was funded by the 3-year head and neck cancer program of the Medical Faculty OWL at Bielefeld University.

Ministerio de Ciencia e Innovación/AEI: Agencia Estatal de Investigación/ 10.13039/501100011033/ y financiado por la Unión Europea “NextGenerationEU”/ PRTR (PID2020-115112RB-I00)

## Availability of data and materials

All data not contained in this manuscript will be deposited in a publicly accessible database upon acceptance.

## Notes

### Competing Interest Statement

The authors have declared no competing interest.

