## Supplementary Figures S1-S4 for "HPV-positive head and neck squamous cell carcinoma cells lose viability during triggered myocyte lineage differentiation"

### SUPPLEMENTARY MATERIAL

**Supplementary Figure S1: Wound healing assay of HPV+ differentiation models.**

**Supplementary Figure S2: HPV+ cells in HNSCC tissue express muscle markers.**

**Supplementary Figure S3: Myocyte-like HPV+ cells in HNSCC tissue display signs of differentiation.**

**Supplementary Figure S4: HPV+ HNSCC cell differentiation induces a loss of Ki67 expression.**

### Supplementary Figures

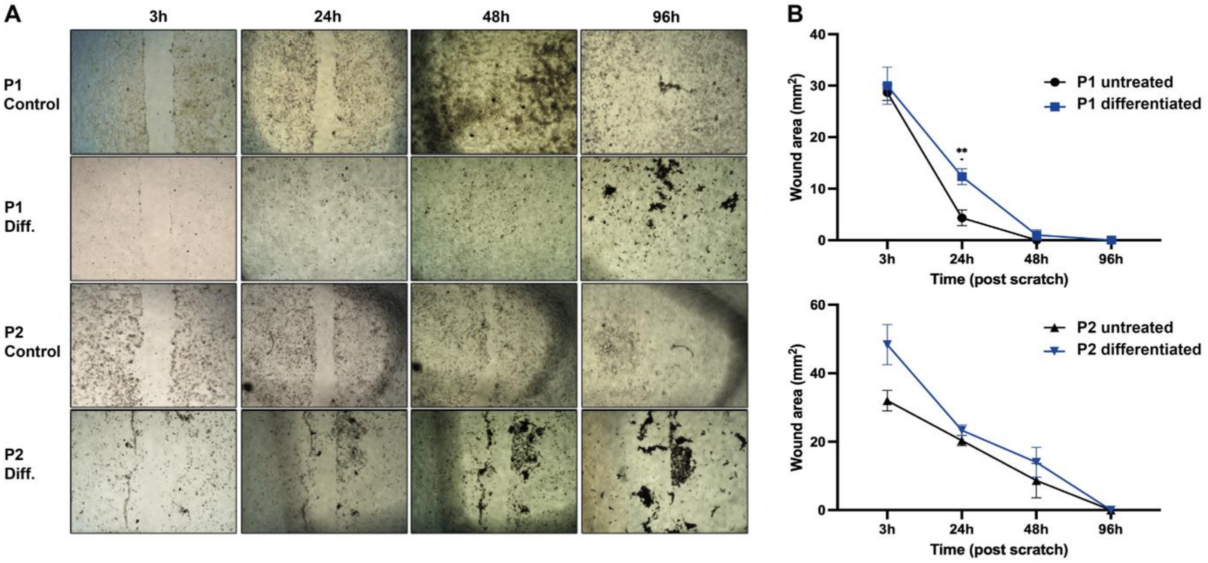


**Supplementary Figure S1: Wound healing assay of HPV+ differentiation models.** **(A)** When treated with the differentiation medium, P1 cells significantly lose wound healing attributes (p-value=0.03), whereas cells of P2 did not change. **(B)** Wound healing area after scratch assay of P1 and P2 undifferentiated and differentiated cells; scale bars = 10 µm.

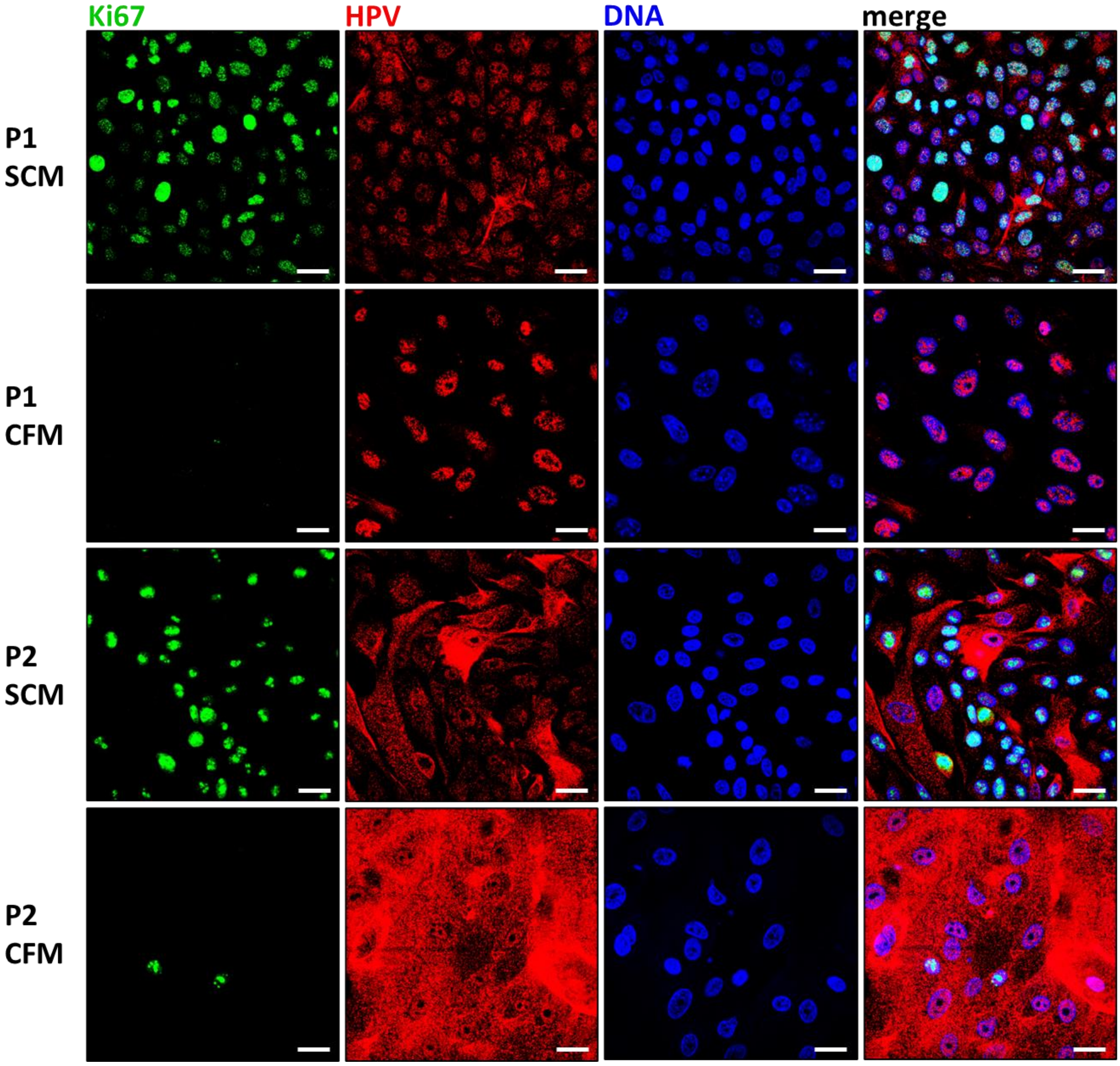


**Supplementary Figure S2: HPV+ HNSCC cell differentiation induces a loss of Ki67 expression.** Cells of P1 and P2 lose Ki67 proliferation marker expression when cultured in differentiation medium (CFM) instead of stem cell medium (SCM). HPV-related proteins are expressed under both conditions; scale bars = 10µm.

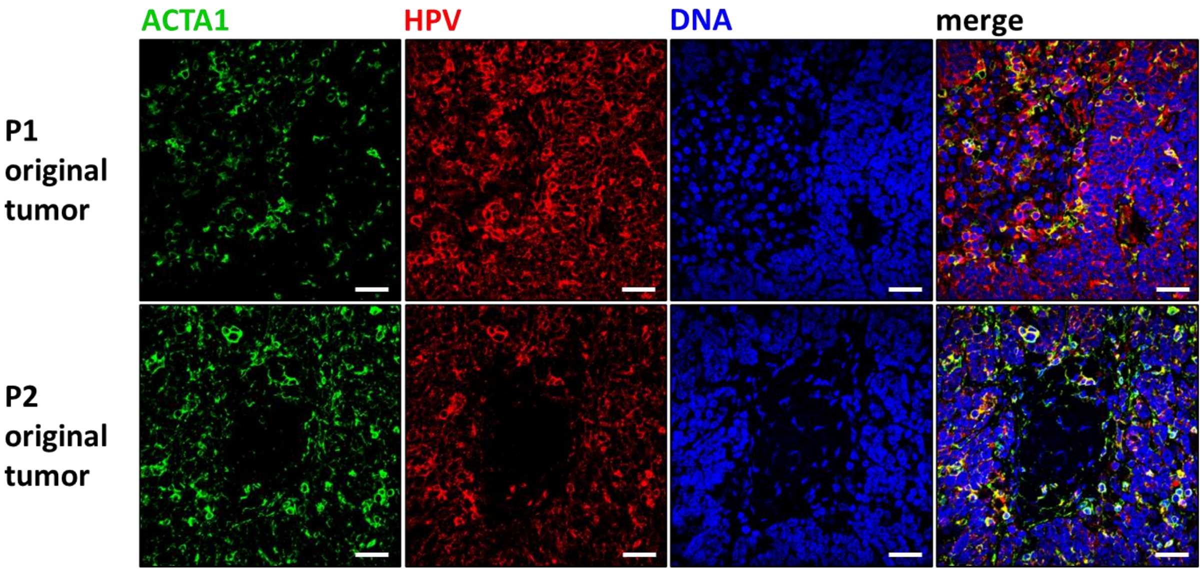


**Supplementary Figure S3: HPV+ cells in HNSCC tissue express muscle markers.** Myocyte lineage protein ACTA1 stains co-positive with HPV-related proteins in human HNSCC tissue of P1 and P2; scale bars = 10µm.

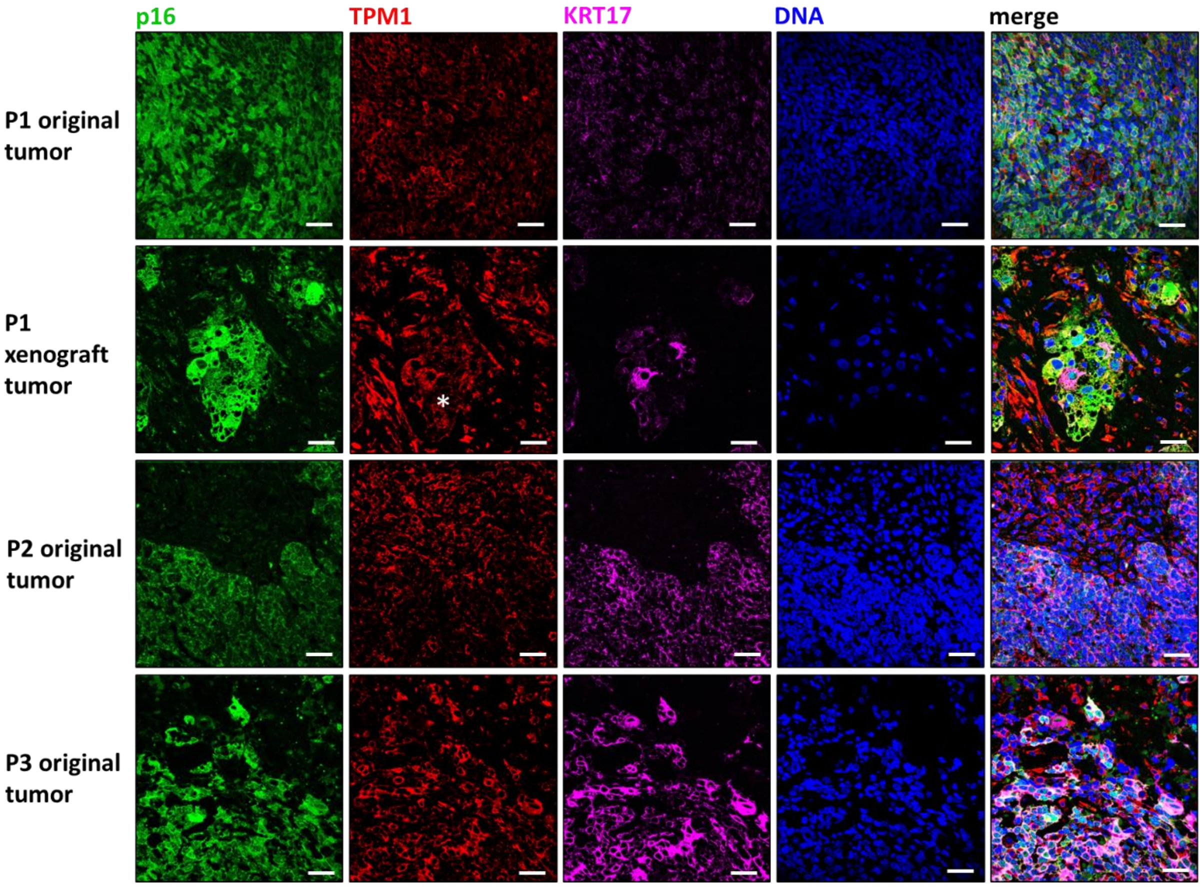


**Supplementary Figure S4: Myocyte-like HPV+ cells in HNSCC tissue display signs of differentiation.** Cells in tumors of P1-P3 show simultaneous expression of HPV surrogate marker p16, myocyte lineage marker TPM1, and differentiation marker KRT17 (Quelle); scale bars = 10µm.
